# Locus Coeruleus tracking of prediction errors optimises cognitive flexibility: an Active Inference model

**DOI:** 10.1101/340620

**Authors:** Anna C Sales, Karl J. Friston, Matthew W. Jones, Anthony E. Pickering, Rosalyn J. Moran

## Abstract

The locus coeruleus (LC) in the pons is the major source of noradrenaline (NA) in the brain. Two modes of LC firing have been associated with distinct cognitive states: changes in tonic rates of firing are correlated with global levels of arousal and behavioural flexibility, whilst phasic LC responses are evoked by salient stimuli. Here, we unify these two modes of firing by modelling the response of the LC as a correlate of a prediction error when inferring states for action planning under Active Inference (AI).

We simulate a classic Go/No-go reward learning task and a three-arm foraging task and show that, if LC activity is considered to reflect the magnitude of high level ‘state-action’ prediction errors, then both tonic and phasic modes of firing are emergent features of belief updating. We also demonstrate that when contingencies change, AI agents can update their internal models more quickly by feeding back this state-action prediction error – reflected in LC firing and noradrenaline release – to optimise learning rate, enabling large adjustments over short timescales. We propose that such prediction errors are mediated by cortico-LC connections, whilst ascending input from LC to cortex modulates belief updating in anterior cingulate cortex (ACC).

In short, we characterise the LC/ NA system within a general theory of brain function. In doing so, we show that contrasting, behaviour-dependent firing patterns are an emergent property of the LC’s crucial role in translating prediction errors into an optimal mediation between plasticity and stability.

**Author Summary:** The brain uses sensory information to build internal models and make predictions about the world. When errors of prediction occur, models must be updated to ensure desired outcomes are still achieved. Neuromodulator chemicals provide a possible pathway for triggering such changes in brain state. One such neuromodulator, noradrenaline, originates predominantly from a cluster of neurons in the brainstem – the locus coeruleus (LC) – and plays a key role in behaviour, for instance, in determining the balance between exploiting or exploring the environment.

Here we use Active Inference (AI), a mathematical model of perception and action, to formally describe LC function. We propose that LC activity is triggered by errors in prediction and that the subsequent release of noradrenaline alters the rate of learning about the environment. Biologically, this describes an LC-cortex feedback loop promoting behavioural flexibility in times of uncertainty. We model LC output as a simulated animal performs two tasks known to elicit archetypal responses. We find that experimentally observed ‘phasic’ and ‘tonic’ patterns of LC activity emerge naturally, and that modulation of learning rates improves task performance. This provides a simple, unified computational account of noradrenergic computational function within a general model of behaviour.

## Introduction

The locus coeruleus (LC) is the major source of noradrenaline (NA) in the brain, projecting to most territories from the frontal cortex to the distal spinal cord. Changes in LC firing have been associated with behavioural changes, most notably the switch from ‘exploiting’ to ‘exploring’ the environment, and the facilitation of appropriate responses to salient stimuli (1,2).

Tonic LC activity is correlated with global levels of arousal and behavioural flexibility, where firing rates increase with rising levels of alertness (1). At the extreme, high rates of tonic firing have been causally related to behavioural variability and stochastic decision making (3). This ‘tonic mode’ has previously been modelled as a response to factors such as declining utility in a task (4) or ‘unexpected uncertainties’ (5), triggering behavioural variability and a switch from ‘exploiting’ a known resource to ‘exploring’ for a new resource.

The LC also fires in short, high frequency bursts. Such phasic activity occurs in animals in response to behaviourally relevant salient stimuli (1,6–8). This phasic response has been described as a ‘network interrupt’ or ‘reset’, which facilitates a shift to shorter-term behavioural planning (9,10). Activating stimuli are those which have an established behavioural significance; for instance, signalling the location of food or the presence of a predator. They may also include stimuli that are highly unexpected (1,11) – although the phasic response will habituate rapidly to novelty alone in the absence of behavioural salience (12).

A series of studies has provided evidence of further nuance to phasic LC responses. Similar to the well-known dopaminergic response, as an animal learns a cue-reward relationship, phasic LC responses will transfer from temporal alignment with an unconditioned stimuli (US) to a predictive, conditioned stimuli (CS+)(1,13). Additionally, rarer stimuli, or those predicting a large reward, elicit a stronger LC response (6,8). In contrast if predictive cues are delivered consecutively, the size of the response appears to decrease (6). The rich array of factors affecting the nature of the phasic response suggests that LC activation is linked to both facilitation of behavioural response and to internal representations of uncertainties and probabilities.

Despite the increasing body of knowledge about the impact of the LC on behaviour, a comprehensive computational account remains elusive – in contrast to the more developed account of other neuromodulators; most notably dopamine, which has been interpreted as a signal of reward prediction error. In particular, existing modelling approaches have generally tackled the tonic and phasic firing responses of the LC as separate modes with distinct functional significance, triggered by different circumstances (4,5,9,10).

Here, we propose that a critical computational role of the LC-NA system is to react to high level ‘state-action’ prediction errors upstream of the LC and cause appropriate flexibility in belief updating via feedback projections to cortex. In brief, our account of noradrenergic activity is based on the fact that the degree of belief updating reflects volatility in the environment and can therefore inform the optimal rate of evidence accumulation and plasticity. The ‘state-action’ prediction error considered in this work is the ‘Bayesian surprise’ or change in probabilistic beliefs before and after observing some outcome. We develop these ideas as neural correlates of discrete updates and action planning under the formalism of Active Inference (AI). AI offers an effective mathematical framework for such modelling, unifying inferences on states and action planning and providing a detailed description of beliefs at each step of a behavioural task (14–17). In taking this formal approach, our description of the LC is integrated into a general theory of the brain function and uses constructs that underwrite the normal cycle of perceptual inference and action selection. This contrasts with previous LC modelling approaches, which have invoked the monitoring of statistical quantities (such as unexpected uncertainty) *per se*.

In the following we apply AI to simulate the updating of beliefs about states of the world – and actions - as a synthetic agent engages with two scenarios (a Go/No-go task with reversal and a foraging task) that elicit archetypal LC responses. Using this approach, we show that the ‘state-action prediction error’ offers an effective predictor of LC firing over both long (tonic) and short (phasic) timescales, without the need to invoke switches between distinct modes. Furthermore, we described how the signal may be broadcast back to cortex to affect appropriate updates to internal models of the environment. This links the error via the LC to model flexibility – bringing two key concepts of the LC together: ‘explore-exploit’ and ‘network reset’. It also produces behavioural changes that agree with experimental knowledge of animal behaviours under noradrenergic manipulation. Finally, the simulations produce realistic LC firing patterns that could, in principle, be used to model empirical responses.

## Methods and Modelling

### Brief overview of Active Inference

Active Inference is a theory of behaviour that has previously been mapped to putative neural implementations (14). The basic premise of AI is that to stay in states compatible with survival, an agent must create and update a generative model of the world (14,18,19). To do this effectively the agent represents the true structure of the world with an internal model that is a good approximation of how its sensations are generated. (Note that in this paper, we often use the term ‘model’ to refer to the agent’s beliefs about states and actions in the world. Technically, these beliefs are posterior probability distributions, which require a generative model to exist.)

The generative model encompasses a set of discrete states and transition patterns that probabilistically capture all the agent’s beliefs about the world and likely outcomes under different actions. The model is formulated as a Partially Observable Markov Decision Process (POMDP), under which the agent must infer its current state, make predictions about the outcome of actions in the future and make postdictions about the landscape it has just traversed. In this context the word ‘state’ refers to a combination of features relevant to the agent, including its location and the cognitive context of that location; i.e., states of the world that matter for its behaviour.

To optimise this model, the agent constantly seeks to minimise variational free energy. This free energy is a mathematical proxy for the difference between the agent’s generative model and a ‘perfect’ or ‘true’ model of the world, and thus must be continually updated for the agent to survive. Estimates of the free energy can be obtained over time by comparing predictions from the generative model with the results of actions in the real world, for instance, by checking whether an action produces the expected sensory feedback. Using this information from the real world, the agent can minimise free energy by moving to expected states or by adjusting the parameters of the generative model itself. The latter allows the agent to optimise the model and/or change its current action plans. Updating proceeds in cycles, with each round of model updates accompanied by predictions that are then checked by selecting and executing an action – in turn allowing a new round of updates (Box 1).

#### Box 1.

##### A quasi-mathematical description of the framework of Active Inference (based on (14))

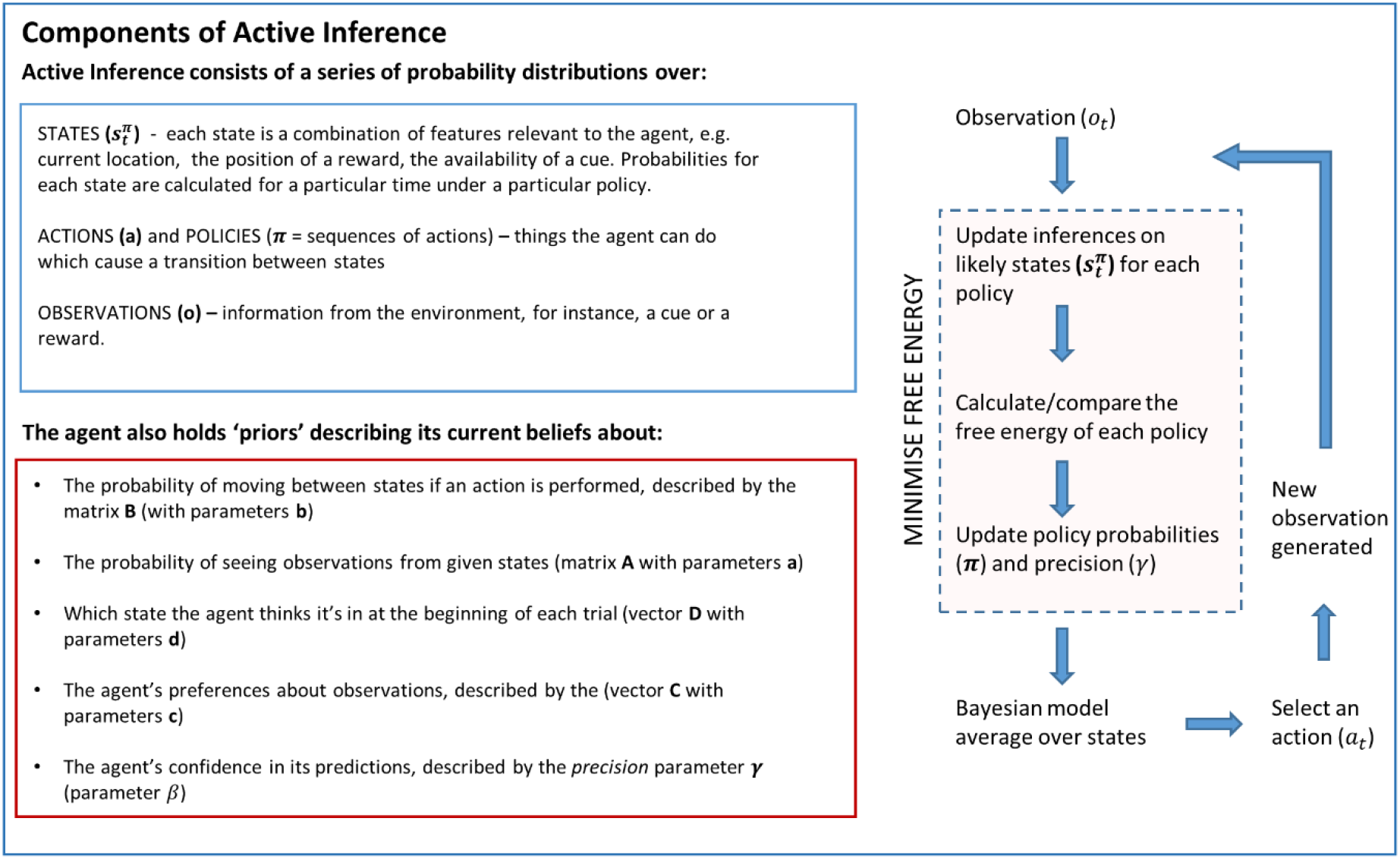

This framework means that each round of updates combines perceptual inference with action selection. Mathematically, updates take the form of a series of iterative updates to parameters that are repeated until convergence (Box 2). It is this machinery that we will map to LC/NA firing and function.

#### Box 2.

##### Five step mathematical outline of the framework of Active Inference

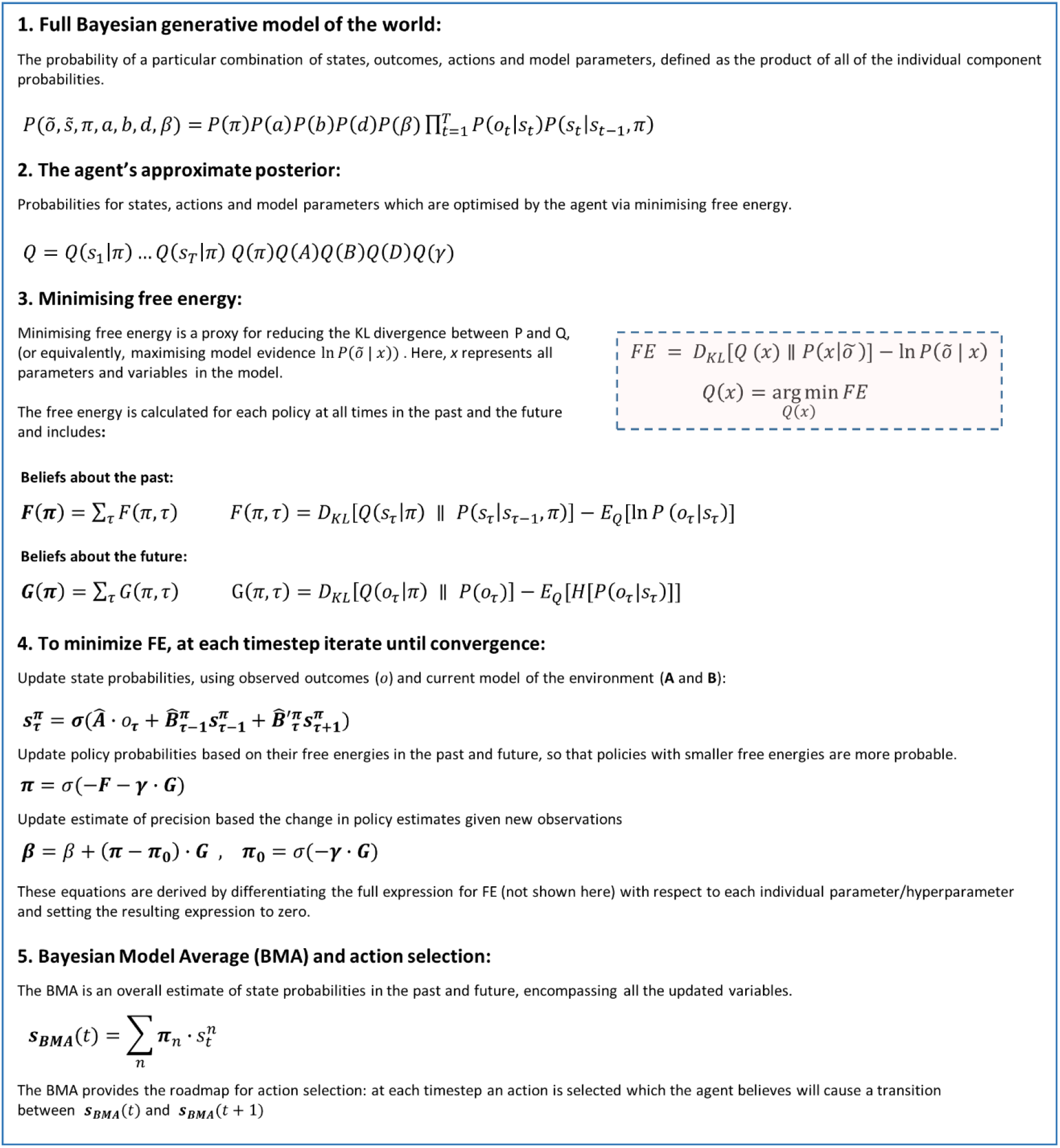

There are two more subtleties that should be noted in this brief description. Firstly, action selection is driven by twin goals – the future attainment of states that the agent holds valuable (utility), as well as the attainment of information when performing an action (epistemic value). Formally, these describe the path integral of free energy expected under competing policies. Thus, agents that act to minimise free energy will end up where they hoped to, while resolving uncertainty about their environment. If policies do not differ in their ability to resolve uncertainty (i.e. no policy will harvest more information) then utility will drive policy selection. It has already been established that this particular cost function explores and exploits in a predictable and mathematically well-defined manner, depending on the relative utility of outcomes and on the uncertainty with which the agent views its environment (15–17,20).

The second important component is the timespan covered by inferences. The agent continually updates its understanding of the past, the present and the future. This means that observations in the present can be used to update inferences on states that occurred in the past – in this way, past events continue to be useful for belief updating long after they occurred. This is just a formalisation of our ability to postdict (e.g., “I started in this context, even if I didn’t know at the time”). Equally, the agent’s knowledge of the world is used to form predictions at future times (e.g., “These are the outcomes I expect under this policy”). The agent not only attempts to use events that have already happened to minimise free energy, but also tries to select actions and inferences which it believes will minimise free energy of future observations.

### A Bayesian Model Average drives action selection

As outlined in Boxes 1 and 2, the generative model comprises probability distributions over states, sequences of actions, precision (confidence in predictions) and observations. The agent also holds prior beliefs about the way these variables interact, for instance, the probability that a particular state will result in a specific observable outcome. At each time step, the agent updates its beliefs about these probability distributions over states, actions and precision by minimising free energy.

Once all updates have been completed the agent combines all of its inferences to produce a Bayesian Model Average (BMA) of states under possible actions. This can be considered as a summary of everything the agent knows about its place in the world – an overall ‘map’ of the states it believes it occupied in the past, the state it occupies now and the states it believes it will occupy in the future. The distribution implicitly includes action planning that is informed by inferences about events in the past. These probabilities can be represented as a ‘state-action heatmap’ showing how the likelihood of different states evolves over time as evidence accumulates and beliefs are updated (see Figure 1). The Bayesian Model Average is then used by the agent to select an action, generating an observation which forms the basis of the next cycle of updates.

### State-action Prediction Errors as a driver of LC activity

Any large change in the state-action heatmap between time steps represents a *state-action prediction error.* These errors indicate that the agent’s beliefs about its past and future states have changed substantially after receiving a fresh observation. Such prediction errors indicate that the agent’s model of the world – including its plan for actions – must change. This may either be because an unexpected stimulus has occurred, requiring an abrupt change in behaviour, or because observations over longer timescales are consistently demonstrating that key components of the model (for example, the observation likelihood (**A**) and state transition (**B**) matrices) are no longer fit for purpose. Crucially, errors originating from both situations are reflected in the state-action prediction error. We propose that they are a driver of LC activity.

The BMA is estimated for each time point and takes the form of a weighted sum over state probabilities (states are weighted by the probability of each policy predicting that state at the given time). To estimate the state-action prediction error during a task, we take the Kullback-Leibler divergence between Bayesian Model Average (BMA) distributions at successive time steps. Mathematically, this reflects the degree of belief updating induced by each new observation. It is often known as a relative entropy, information gain or Bayesian surprise. The following expressions describe the BMA (upper equation) and the state-action prediction error (lower):

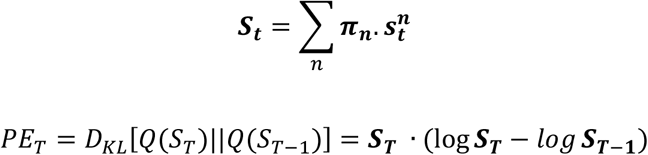

Here, 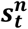 refers to the vector of probabilities of states at time *t* under policy ***π_n_***, whilst *Q* refers to the agent’s current set of beliefs (i.e. *Q*(*s_T_*) indicates the probability distribution for *s_T_* expected under current beliefs).

Prediction errors over shorter timescales (i.e. between actions, during the iterative cycle of belief updating) are an integral feature of AI. The state action prediction error, in contrast, represents a global error: it is expressed over the timescale of a behavioural epoch as a *response* to the outcome of belief updating that precedes action selection.

### LC feedback: flexible model learning promoted by prediction errors

Why might it be useful for the LC to respond to state-action prediction errors? We suggest that one important function is that such errors require a specific modulation of distributed cortical activity encoding representations of the structure of the environment, particularly in frontal cortex. This modulation would boost the flexibility of internal representations (where our matrices would be formed by particularly connected cell assemblies in frontal cortex) and increase their responsiveness to recent observations. In vivo, this may be mediated by the release of noradrenaline from LC projections to the frontal cortex occurring in response to prediction errors.

The need for flexible model updating is directly relevant to a related challenge for Active Inference models; namely, the rate at which the agent’s experience is assimilated into its model. Addressing this issue provides a pathway for modelling the effect of LC activation and closes the feedback loop between brainstem and cortex. So what computational role does NA have in facilitating adaptive flexibility?

Under AI, the agent’s model of the world is encoded by a set of probability distributions that keep track of the mappings between states and outcomes, and between states occupied at sequential time points. These mappings are encoded by Dirichlet distributions, the parameters of which are incremented with each instance of a particular mapping the agent experiences (illustrated in Figure 5) (14,20). However, difficulties arise when environmental contingencies change, because the gradual accumulation of concentration parameters is essentially unlimited. Accumulated experience can come to dominate the agent’s model, with new information having little effect on the agent’s decisions. This occurs because the generative model does not allow for fluctuations in probability transitions, i.e. environmental volatility. This issue can be finessed by adding a volatility or decay factor *(a),* which effectively endows the generative model with the capacity to ‘forget’ experiences in the past that are not relevant if environmental contingencies change (as per code available from http://www.fil.ion.ucl.ac.uk/spm/ (14)).

**Figure 5.**
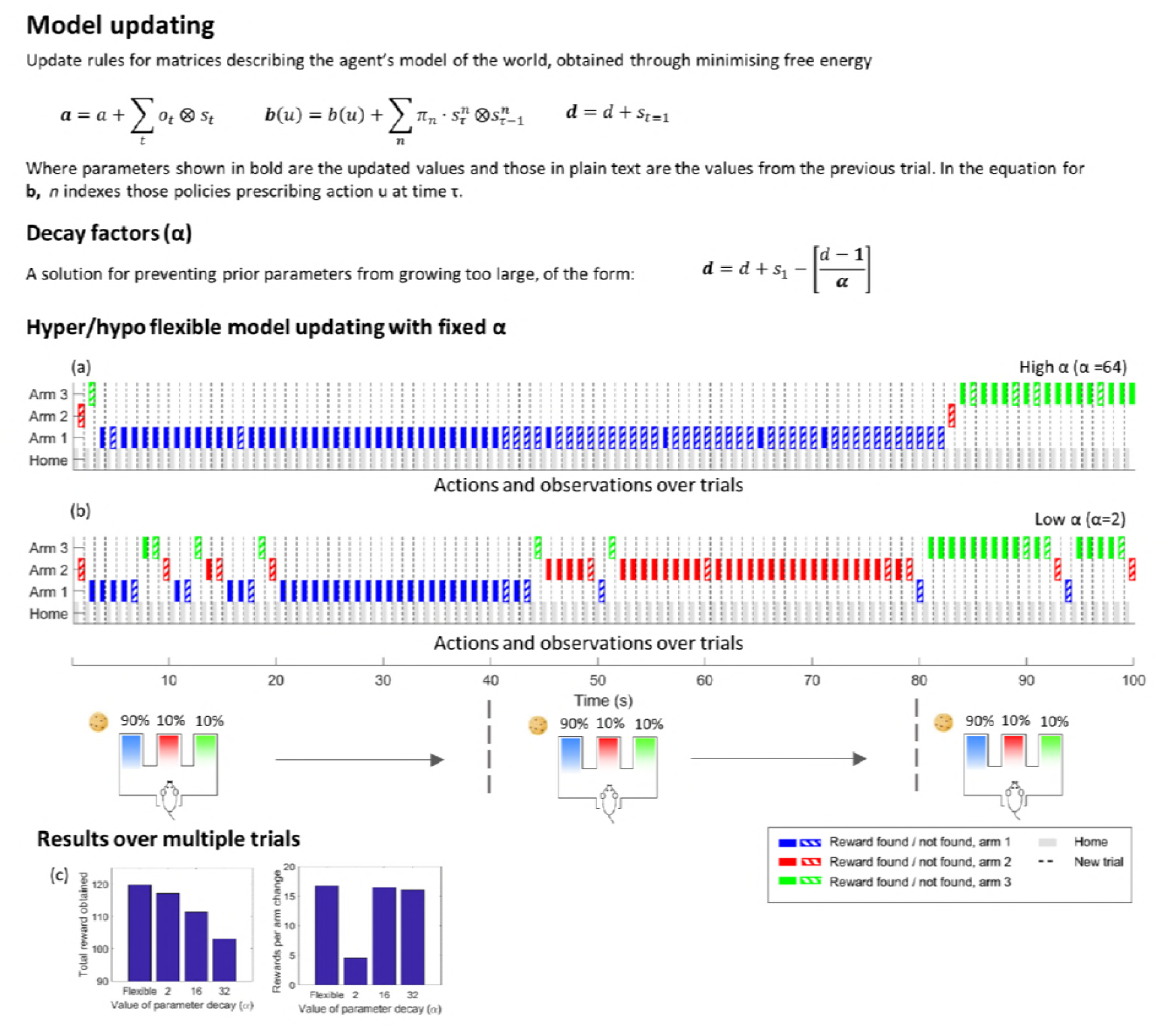
Details of update rules and comparison between flexible and fixed parameter decay. Upper panel: rules for updating Dirichlet parameters. Each parameter is incremented every time a certain mapping is observed. Lower panel (a), (b): examples of agents in the foraging task with values of the decay parameter set high or low. When α is too high, the agent is inflexible and fails to respond to the altered reward probability distributions despite consistently failing to obtain a reward. When α is low, the agent is hyperflexible and often visits a new location after a single unrewarded choice. (c) Mean overall rewards and mean ratio of rewards to changes of location when the same task (with a rule change every 40 trials) is played 100 times. The ‘flexible’ agent’ is endowed with a variable α ranging from 2 to 32 along a sigmoid curve. This agent receives more rewards overall and still has the highest ratio of rewards to changes of location when compared with agents given fixed values of α in the same range.

In the context of reversal learning, this is not a trivial adjustment but a crucial addition to the generative model which enables AI agents to adapt flexibly. However, the level at which to set the decay term poses a further challenge: if the decay is too big, the model is too flexible and will be dominated by its most recent experiences (as all the other terms will have decayed). If the decay is too small concentration parameters may accumulate too slowly, rendering the model too stable.

There are several ways one can optimise this ‘forgetting’ in volatility models. One could equip the Markov decision process with a further hierarchical level modelling fluctuations from trial to trial – as in the hierarchical Gaussian filter (21). A simpler (and biologically plausible) solution is to link the decay factor to recent values of state-action prediction error via the LC. In other words, equip the agent with the prior belief that if belief updating is greater than expected, environmental contingencies have become more volatile.

This produces flexibility in model learning when prediction error is high (low α) but maintains model stability when prediction error is low (high α). We have modelled this feedback using a simple logistic function to convert prediction error into a value for *α*:

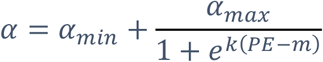

where *PE* is the prediction error seen during the trial (in tasks with more than one prediction error per trial, the maximum error is used), k is a gradient and *m* is a mean (i.e., expected) value. In all simulations presented below, ∝_min_=2, ∝_mix_=32, k=8, and *m* is set as a proportion of the maximum prediction error possible in each task.

Under this scheme, a brief but large prediction error ‘boosts’ the impact of a recent experience upon the agent’s model of the world. This occurs by temporarily increasing the attrition of existing, experience dependent parameters encoding environmental contingencies. Crucially, this causes recent actions and observations to have a greater effect on the Dirichlet distributions than they would otherwise. If prediction errors then decrease, the model stabilises again. However, if actions consistently produce large prediction errors then the underlying model parameters will gradually lose their structure – equivalent to the flattening of probability distributions that form the agent’s model – leading to greater variability in action selection.

## Results

The simulations reported in this paper suggest that behavioural contexts that produce large state-action prediction errors are also those that produce archetypal LC responses in experimental environments. Below, we describe the emergence of phasic and tonic activity in two tasks, as a response to changes in prediction error. We initially present results without the LC feedback in place before showing how both simulations are improved by modelling the LC as a link between prediction errors and model decay / volatility.

### Go/No-go task

A simple ‘Go-No-Go’ game modelled under AI is shown in Figure 1 (using MATLAB code based on (14)). In this game, the agent (depicted as a rat) starts in a ‘ready’ state – location 1 – and must move to location 2 to receive a cue. When the cue is received the agent may either move back to location 1 or seek a reward at location 3. The agent has a strong preference for receiving the reward but an aversion to moving to location 3 and remaining unrewarded. This is represented in the game by a notional ramp which forces the agent to expend physical effort in seeking the reward. There are six available states, which between them describe the different combinations of features relevant to the agent during the game. Learning is mediated through updates to the A and D matrices, which encode likelihood mappings between hidden states of the world and outcomes – and prior beliefs about initial states.

**Figure 1.**
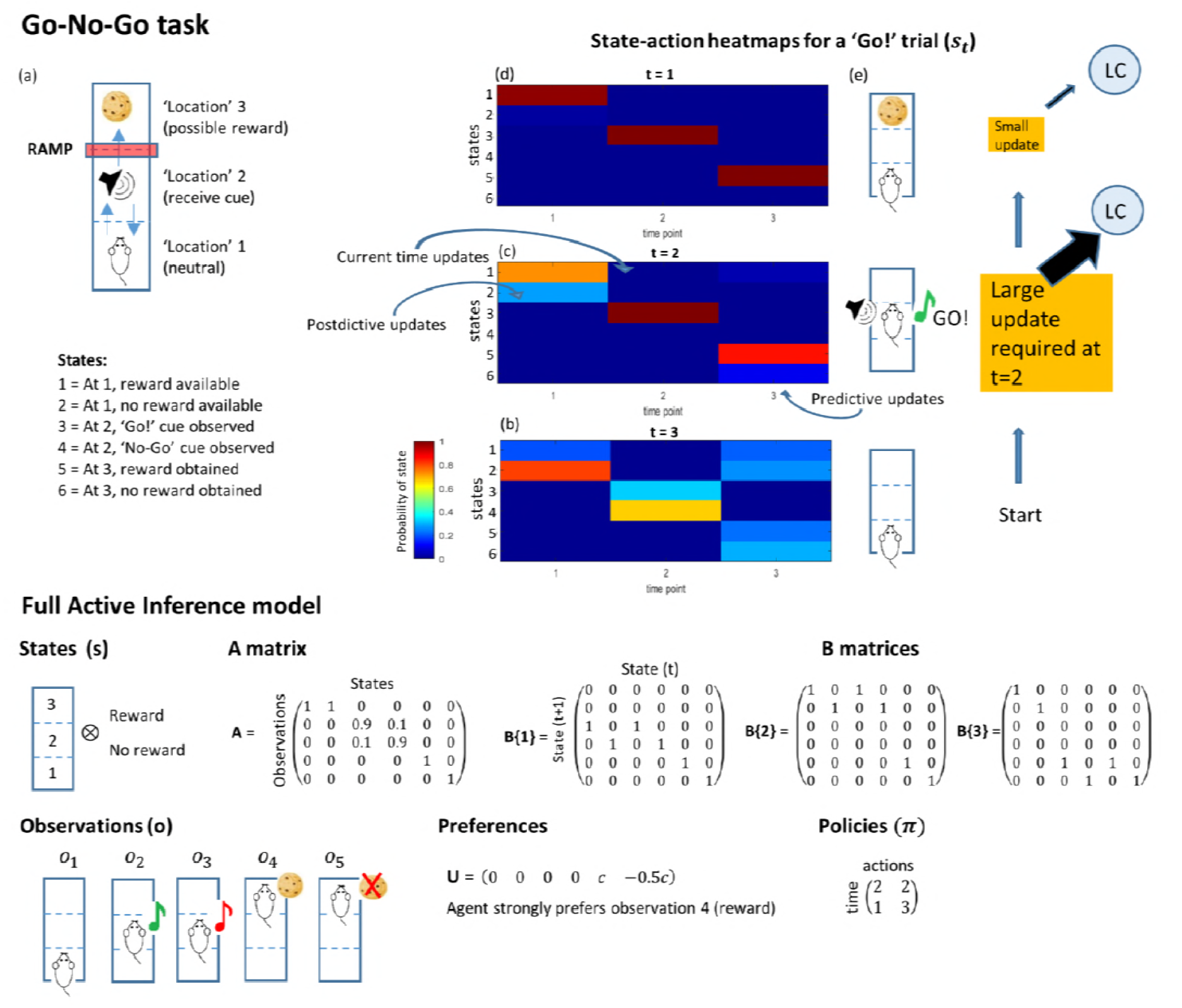
Simple ‘Go-No-Go’ game modelled under AI. (a) Structure of the task (see main text) (b)-(d) The state-action heatmap showing inferences on the agent’s state over a rare ‘Go!’ trial. Large updates are required at t=2, after the animal receives the ‘Go’ cue which forces it to update its action plans and state inferences. This update is proposed to cause a large, time specific input into LC (e), which causes a sudden phasic burst of LC activity. The lower part of the figure shows the full modelling of the Go-No-Go task, with components as described in Box 1.

At each time point, the agent’s beliefs are summarised in the Bayesian Model Average, represented graphically as a state-action heatmap. Figure 1(b) shows a representation of the agent’s beliefs about states at the beginning of a new trial in which the ‘Go’ cue is heard. The agent is ‘well trained’; that is, it has an accurate understanding of the relationship between the cue and the availability of the reward, and of the fact that the ‘Go’ cue is rare (here, the cue probability is 10%). In our modelling, we trained the synthetic rat by running the simulation for 750 trials. We then used the learnt priors as the starting point for the ‘well trained’ case.

Given its knowledge of the game, the agent begins with a strong belief that it is beginning the trial in state 2 (in which a reward will not be available). It also makes predictions for the states it believes it will occupy later in the trial: at t=2, it believes it is likely to occupy state 4 – corresponding to the occurrence of the ‘No-go’ cue, but also entertains a slight possibility that the ‘Go’ cue might still appear. The agent is much less certain in its predictions for t=3, but still holds a higher probability that it will end up in one of the unrewarded end states.

At the next time point (at t=2, Fig. 1(c)), the agent updates its state-action heatmap, making new inferences on the probabilities of different states in the past, present and future, based on its most recent observations. If it has received the rare ‘Go’ cue, it will have to update its predictions for its state at the end of the game, in addition to altering its inferences about the state in which it started at t=1 (a process of postdiction about past states based on new information). The agent therefore has to make a large, sudden update to its BMA heatmap at t=2. By t=3 (Fig. 1(d)), the agent has received the reward as predicted, and knows with certainty where it is and where it has been. Only small updates are required to its estimates at this point.

Simulated prediction errors during this task are shown in Figure 2, in which LC firing is modelled by converting the prediction error to a firing probability via a sigmoid activation function. In this simulation the prediction error does not modulate learning and the decay parameter *α* has been set to a fixed value. During the task, an agent who is well trained shows large peaks of state-action prediction error when the reward-predicting cue is presented, resulting in phasic activity in the LC as seen experimentally (6,22). The underlying reason for this error is a large, quick shift in action planning, from the (more likely) ‘No-go’ outcome to the rare ‘Go’ situation.

**Figure 2.**
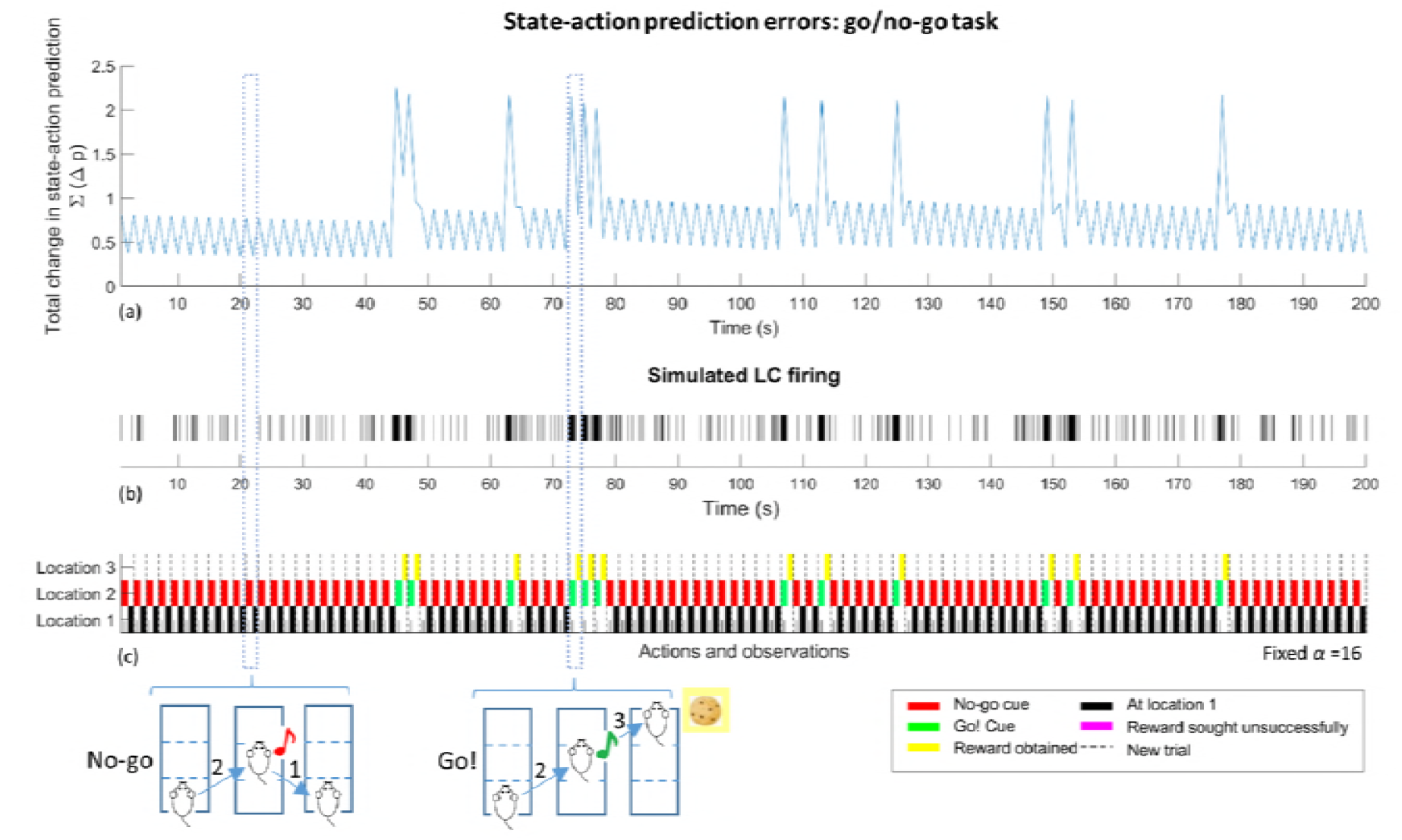
Plot of prediction error (a), simulated LC spiking (b) and behaviour (c) during 100 trials of the Go/No-Go task described in main text. In (a) the raw prediction error is extracted for t=2, when the animal receives a cue (this is the error between t=1 and t=2) and t=3 when the animal receives feedback on its response to the cue (the error between t=2 and t=3). Because the prediction error explicitly evaluates differences between update cycles, there is no error available for the first time point. Each trial has therefore been collapsed to two time points, each lasting 1 second. In (a) the occurrence of the ‘Go’ cue causes strong peaks in prediction error. This is converted into a simulated LC firing rate in (b), showing phasic LC activation when the ‘go’ cue is heard. Plot (c) is a graphical representation of behaviour during the task at times t=2 and t=3.

### Foraging

To supplement the above go no-go task we modelled a foraging task, depicted in Figure 3. On every trial in this task the agent searches for a reward in one of three arms. In one arm, the probability of finding a reward is high (90%), whilst in the others the probability is low (10%). The probabilities are held constant for a set number of trials, during which time the agent accumulates beliefs about the likelihood of finding a reward in each location. Typically, once the agent has been rewarded in one location numerous times it will build a strong prior probability on the availability of a reward in that location (reflected in updates to elements of the **B** matrix). In the example shown in Figure 3 the agent begins by exploring the arms until it has seen a reward in arm 1, after which it continues to visit this location. After a set number of trials, the location of the high probability arm is shifted. When this happens, the agent’s established model of the world no longer provides an accurate explanation of its experiences. As expected rewards fail to materialise, state-action prediction errors arise. Under our model, this causes an increasing tonic rate of LC activity whilst new priors are learnt and behaviour changes.

**Figure 3.**
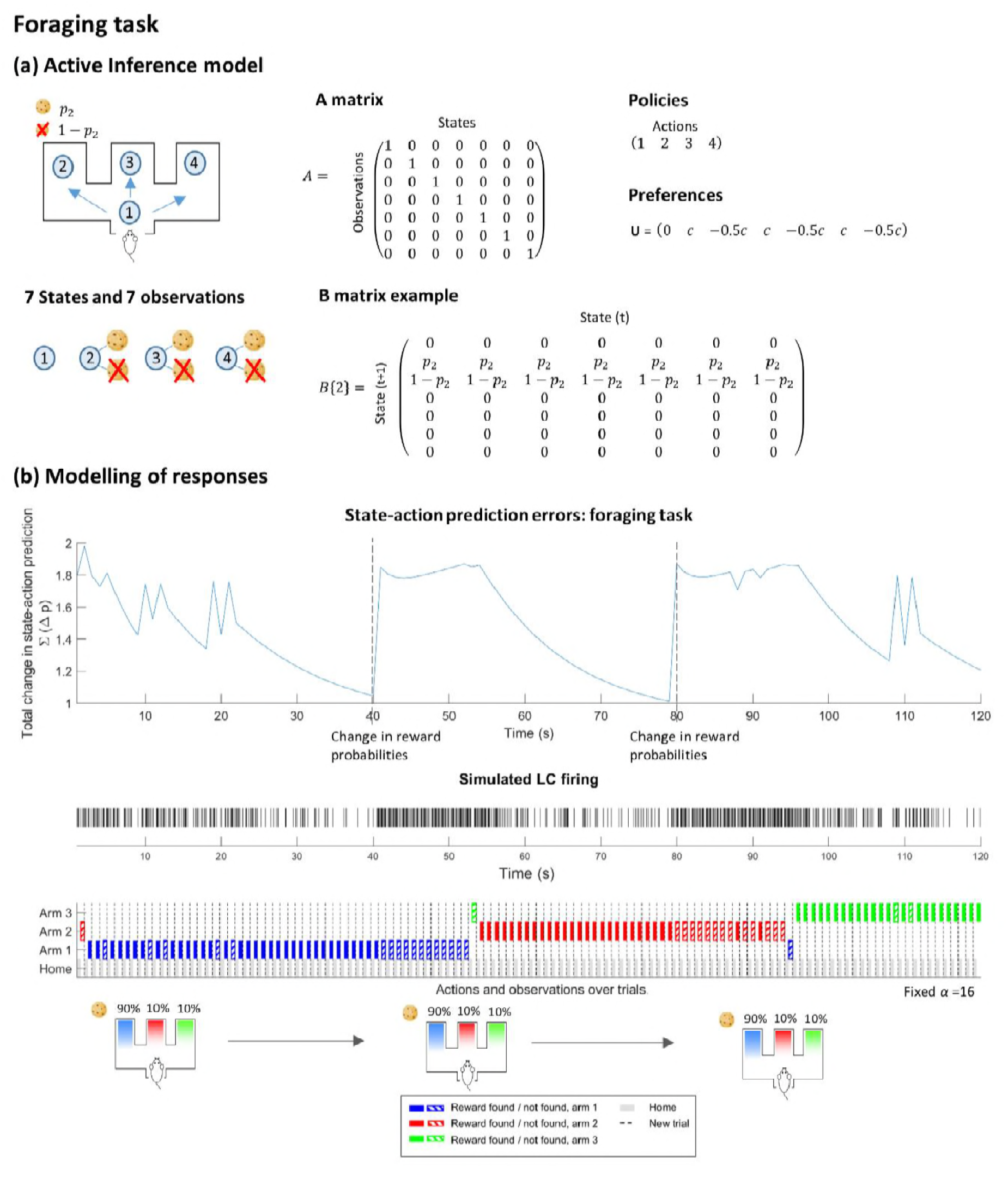
Modelling of a 3-arm foraging task under Active Inference. Upper plot: the mathematical structure of the task. There are seven states, including one neutral starting point and 3 arm locations which can be combined with either a reward / no reward. There are 7 observations; here these have a 1-to-1 mapping to states (A matrix). Actions 1-4 simply move the agent to locations 1-4 respectively. The probability of obtaining a reward in a given arm (p_2_ for action 2, above) is held static for a fixed number of trials, with one arm granting a reward with a 90% probability and the others with 10% probability. This is then switched, so that the agent must adjust its priors and its behaviour. Lower plot: State action prediction errors and LC responses over a typical run of 100 trials.

### Flexibility in model learning: closing the loop

When the full feedback loop between prediction error and model decay is introduced, there are improvements in performance in the simulations of both the Go/No-go and Foraging tasks (Figure 4).

**Figure 4.**
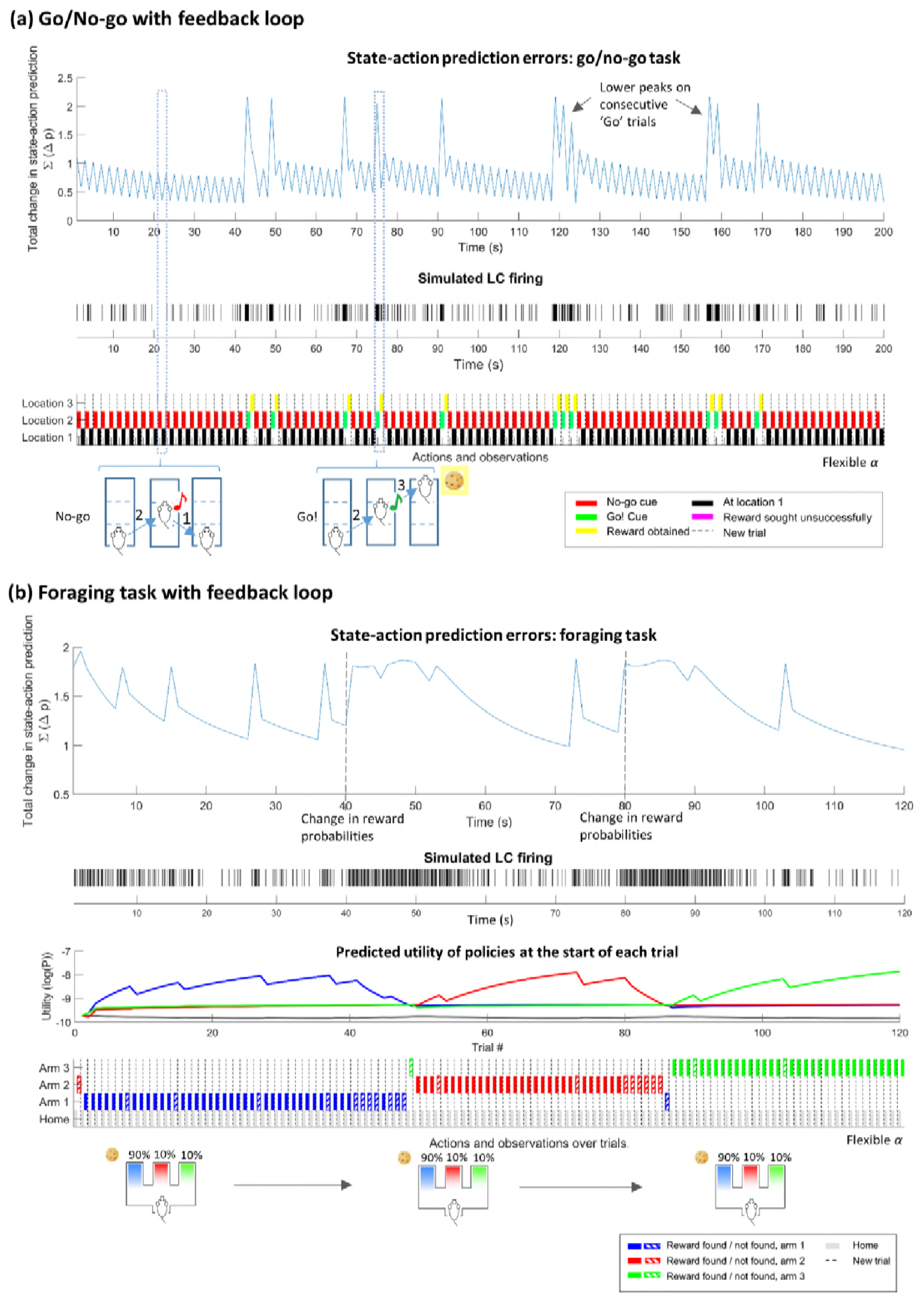
Application of the feedback loop between state-action prediction error and parameter decay to the Go/No-go (a) and Foraging tasks (b). See main text for description.

One consequence during the Go/No-go game is that multiple consecutive ‘go’ trials produce clearly reduced peak heights (as has been recorded in terms of LC activation in the same context (6)). This is due to the continual modulation of the agent’s prior beliefs about whether each trial will be a Go or No-Go context (encoded by *d* parameters that accumulate experience about initial states). With a brief high prediction error, the update prioritises the recent experiences of the agent: after a few consecutive ‘Go’ trials this creates a distribution with a higher probability of the ‘Go’ context than would be suggested by the statistics of the rat’s entire experience in the game.

In the foraging task, the dynamic modulation of model building allows prediction errors to reduce more quickly when the rat is settled into the ‘exploit’ mode of harvesting a reward in a reliable location, promoting model stability (Figure 4b). When the reward is no longer available, errors mount and the increase in model decay causes the agent to make more explorative choices. This contrasts with the same task simulated with fixed values of *α* (Figure 5): when the model is hyper-flexible, the agent often switches behavioural strategy after a single failed trial; when the model is inflexible, the agent takes a large number of trials to visit a new location. Over multiple trials, the agent with a dynamically varying *α* consistently secures more rewards than agents with fixed α values taken from the same range (Figure 5c).

Finally, the application of this scheme to a reversal learning scenario under the Go/No-Go game is described in Figure 6. As expected, the well-trained agent begins the session by showing a phasic response in prediction error / LC firing in response to the ‘Go’ cue (cue 1). At trial 35, the meaning of the two cues switches so that the ‘Go’ context is predicted by cue 2. At the reversal, state-action prediction errors cannot be resolved and LC firing switches to a higher tonic level. During this period, model updating – and behaviour – becomes more flexible and the new rules of the task are learnt. Eventually the high levels of tonic activity fall away and phasic responses to the new ‘Go’ cue re-emerge; coupled with a lower level of tonic activity. This mirrors the pattern of LC firing recorded in monkeys during the same task (22).

**Figure 6.**
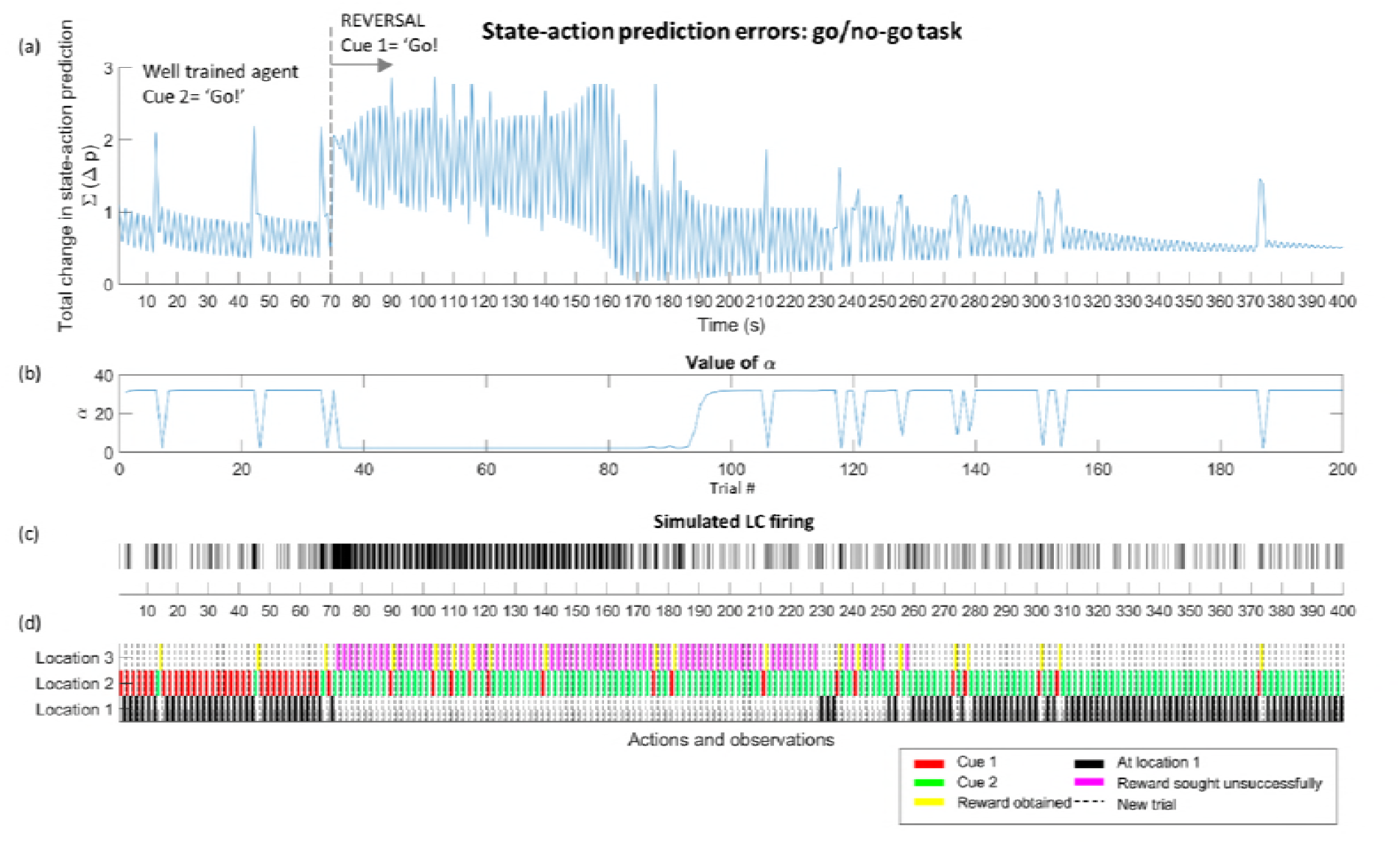
Reversal learning during the Go-No-Go game. The agent begins with a well-trained understanding (via 750 trials of training) that cue 2 indicates that a reward is available. At trial 35 (t=70) the cue/context relationship is reversed, and the agent must now learn that cue 1 indicates the ‘Go’ context. This initially causes numerous unsuccessful trials, violating the learnt model and producing high prediction errors (a). Note that prediction errors are initially elevated at both timepoints in each trial because both the previously rare cue and the subsequent lack of reward are unexpected. These prediction errors result in a lowering in the parameter decay factor (b), which in turn flattens the agent’s priors causing more variability in behaviour. Eventually the agent learns the new contingencies and the model stabilises, with the reemergence of phasic bursts of LC activity on ‘Go’ trials (a, c). From trial 125 onwards, the peak of phasic activity begins to transition towards the presentation of the cue rather than the reward. This is also seen during the training period of the well-trained agent shown in Figure 2 and 4(a).

## Discussion

We propose that the LC fulfils a crucial role, linking prediction errors (or Bayesian surprise) during the planning of actions to model decay – a form of learning rate. Using this approach, we have reproduced the following experimentally observed LC characteristics:

- Phasic responses during a Go/No-Go paradigm such as the one described in (6,22). Here, cues predicting a reward (for which the animal must perform an action) elicit clear phasic LC responses, which stand out against a background of lower overall tonic activity.
- Consecutive rare stimuli (‘Go’ trials) result in progressive reductions in LC phasic response
- Responses during a reversal of contingencies in the Go / No-go task as described in (22), during which phasic responses are lost in favour of higher tonic activity during the reversal. This is thought to allow behavioural flexibility, which in turn allows the learning of new contingencies (reviewed in (4)) As new rules are learnt, phasic responses eventually re-emerge on the presentation of the new reward-predicting cue.
- A more general link between the ‘exploration’ mode of behaviour and higher tonic levels of activity. Whilst direct measurements of LC activity during explore-exploit paradigms are lacking, the link is strongly suggested by indirect experimental evidence. For instance, Tervo et al (3) demonstrated highly variable behavioural choices in rats when the activity of LC units projecting to ACC was held artificially high via optogenetic manipulation. Other studies have also demonstrated (23,24) that an increase in pupil size (a correlate of LC activity) occurs in parallel with behavioural flexibility and task disengagement.

### Neurobiology

In previous Active Inference literature the calculation of Bayesian Model Averages has been mapped to the dorsal prefrontal cortex (14). This is one of the frontal regions known to send projections to LC (25,26) and is a candidate for the calculation of state-action prediction error (although we accept that without further experimental work such anatomical attributions are largely speculative). Experimental evidence for a neural representation of a distinct prediction error based on states, rather than rewards, has also been found in dorsal regions of the frontal cortex in a human MRI study (27).

Turning to the LC-prefrontal connections and the modulation of model updating, converging experimental evidence suggests that working models of the environment are reflected by ACC activity. Activity in the ACC has been shown to correlate to many factors relevant to the maintenance of a generative model, including reward magnitude and probability (for review see (28)), estimation of the value of action sequences and subsequent prediction errors (29,30) and the value of switching behavioural strategies (31). Marked changes in activity in ACC have been observed at times thought to coincide with significant model updating and occur in parallel with explorative behaviour – an event that has been directly linked to increased input from locus coeruleus (3,32). Similarly, a direct ACC/ LC connection has also been found in response to task conflicts (33). ACC activity is also correlated with learning rate during times of volatility, such that when the statistics of the environment change, more recent observations are weighted more heavily in preference to historical information (34). This evidence provides a solid foundation for the hypothesis that the LC modulates learning rate by governing model updating via ACC. Specifically, we propose that the release of noradrenaline would cause a temporary increase in the susceptibility of model-holding networks to new information. At a cellular level, this would lead to NA effectively breaking and reshaping connections amongst cell assemblies.

In vitro investigation of the cellular effects of noradrenaline provides support for this idea, indicating that noradrenaline may suppress intrinsic connectivity of cortical neurons, causing a relative enhancement of afferent input (1,35,36). Sara (37) and Harley (38) also suggest that LC spiking synchronises oscillations at theta and gamma frequencies, allowing effective transfer of information between brain regions during periods of LC activity. This may allow enhanced updating of existing models with more recent observations. A role for the LC in prioritising recent observations during times of environmental volatility has been explicitly suggested experimentally (39) and is supported by evidence regarding the critical role of LC activation in reversal learning, e.g. (40).

We note that if the LC is indeed responding to prediction errors, model updating is likely not the only functionality it has. For instance, LC activation has been experimentally linked to the potentiation of memory formation (37,41,42), analgesic effects (43,44) and changes to sensory perception for stimuli occurring at the time of LC activation (1,45,46). These are all reasonable responses to a large prediction error: the increase in gain on sensory input may ensure that salient stimuli are more easily detectable in the future, whilst enhanced formation of memory might ensure that mappings between salient stimuli and states are remembered over longer timeframes. Similarly, the temporary suppression of pain may facilitate urgent physical responses to important stimuli (for instance, allowing action in response to a stimulus indicating the presence of a predator). The possibility that the LC has the capacity to provide a differentiated response to prediction error is supported by recent work indicating that existence of distinct subunits with preferred targets producing different functional effects (44,47–49).

### Relationship to existing models of LC function

The ideas described above are not a radical departure from existing models of LC function – but use the theory of active inference to integrate similar concepts into a general theory of brain function, without invoking the need for monitoring of ad-hoc statistical quantities.

The adaptive gain theory proposed by Aston Jones and Cohen (4) proposes that the LC responds to ongoing assessments of utility in OFC and ACC by altering the global ‘gain’ of the brain (the responsivity of individual units). Phasic activation produces a widespread increase in gain which enables a more efficient behavioural response following a task-related decision; however, when the utility of a task decreases, the LC switches to a tonic mode which favours task disengagement and a switch from ‘exploit’ to ‘explore’.

The mechanism we have described reproduces many elements of the adaptive gain theory, with the important exception that different LC firing patterns promoting explorative or exploitative behaviour are an emergent property of the model rather than a dichotomy imposed by design. Since the probability assigned to individual policies is explicitly dependent on their utility (in combination with their epistemic value) a large state-action prediction error will ultimately reflect changes in the availability of policies which lead to high utility outcomes. This may be a positive change, as is the case when a cue indicates that a ‘Go’ policy will secure a reward, or a negative change, when rewards are no longer available in the foraging task. This link is demonstrated in Figure 4 for the foraging task, where increases in prediction error / LC firing occur in tandom with abrupt changes in the agent’s assessment of a given policy’s utility. Both the LC response, and the underlying cause (prediction error), show a shift between ‘phasic’ and ‘tonic’ modes (although it is entirely possible that coupling mechanisms within the LC also act to exaggerate the shift and cause the LC to fire in a more starkly bi-modal fashion, as suggested by computational modelling of the LC (4,50)). As described above, a short prediction error will act to heighten the response to a salient cue over the short term, whilst a large, sustained prediction error – occurring in parallel with declining utility in a task – will act to make behaviour more exploratory.

Yu and Dayan have proposed an alternative model where tonic noradrenaline is a signal of ‘unexpected uncertainty’, when large changes in environment produce observations which strongly violate expectations (5). This is described as a ‘global model failure signal’ and leads to enhancement of bottom-up sensory information. We have focused on a similar ‘model failure’ signal which allows larger changes to learning about the structure of the model itself – but using the inferences of states within AI as our driver, avoiding explicit tracking of the statistics of ‘unexpected uncertainty’. Rather, we compute model failure in terms of ‘everyday’ errors in predicted actions and sensations. Our model is also in line with the ‘network reset’ theory proposed by Bouret et al, in which LC phasic activation promotes rapid re-organisation of neural networks to accomplish shifts in behavioural mode (10), see also (9). Large changes in configuration of the state-action heatmap alongside the updates to internal models above would similarly constitute network re-organisations with the result of changing behaviour. Importantly, state-action updates precede action selection, placing LC activation after decision making / classification of stimuli, but before the behavioural response. This order of events is in keeping with experimental evidence showing that LC responses do indeed consistently precede behavioural responses (51,52). This also parallels the ‘neural interrupt’ model of phasic noradrenaline proposed by Dayan and Yu (53), in which uncertainties over states within a task are signalled by phasic bursts of noradrenaline, causing an interrupt signal during which new states can be adopted.

More recently Parr et al have described an alternative active inference-based model of noradrenaline in decision making (54). Under this model, noradrenaline and acetylcholine are related to the precision assigned to beliefs about outcomes and beliefs about state transitions. That is, the agent assigns a different weight to any inferences made using the **A** matrix (modulated by release of acetylcholine) or the **B** matrix (modulated by noradrenaline) in its updates. This approach captures some of the interplay between environmental uncertainty and release of noradrenaline. Our formulation also speaks to these uncertainties – without the need to introduce new volatility parameters, or to segregate cholinergic / noradrenergic response into separate modulators of likelihood and transition (i.e., **A** and **B** matrices). Both approaches target the coding of contingencies in terms of connectivity (i.e., probability matrices). Parr et al consider the optimisation of the precision of contingencies. Conversely, we consider the optimisation of precision from the point of view of optimal learning rates. In other words, the confidence or precision of beliefs about outcomes likelihoods and state transitions can itself be optimised based on inference (about states) or learning (about parameters) in the generative model.

The key contribution of the current work is to link inference to the precision of beliefs about parameters via learning. This addresses the issue of how model parameters are learned and updated and allows an AI agent to make substantial changes to the architecture of its model in times when environmental rules have shifted. The ensuing behaviour produces the archetypal phasic-tonic shifts in LC dynamics, and links LC responses to the outcome of decision on stimuli, as suggested by in-vivo recordings; summaries of which can be found in (4,11).

The difference between these two applications of Active Inference illustrates a broader point about the way in which the theory is used to describe neuromodulation. Current versions of Active Inference have conceived of neuromodulatory systems as reflections of precision, altering the weights assigned to components of the agent’s model *during* a continuous cycle of updates. This underlying modulation is capable of drastically altering the inferences the agent makes about likely states and actions. Here we have offered a different view, in which noradrenaline is proposed to respond to the *outcome* of an update cycle. This enables us to endow an active inference agent with a noradrenergic response which relates activity in the locus coeruleus to the outcome of decisions and to subsequent changes to action planning. These responses are then linked back to changes in the underlying structure of the agent’s model – again outside of the cycle of inferences. Placing such responses above the update cycle moves them closer to the level of action selection and allows us to reproduce many aspects of LC dynamics observed empirically.

## Future work

Once validated through experimental work, models of this type can provide insight into symptoms of disorders which have been linked to LC dysfunction. For example, attention deficit hyperactivity disorder (ADHD), which is characterised by inattention and hyperactivity, has been associated with elevated tonic LC activity (1). Under our model, high tonic firing rates would cause a persistently high ‘model decay’. This would cause similar outcomes to those demonstrated for the hyper-flexible foraging agent (Figure 5), which cannot build a stable structured model of the environment and reacts to even minor violations of predictions by changing its behavioural strategy. Pharmacological interventions which lower tonic LC firing rates may ameliorate symptoms by allowing structured models to emerge, guiding appropriate phasic responses and producing more focused behavioural strategies.

Several lines of future work are available to test components of the prediction error / LC theory. Firstly, a clearer understanding of the drivers of LC responses could be pursued through in-vivo recordings in PFC, ACC and LC. This would help to confirm if calculations of prediction error (or utility/estimation uncertainty, under other theories) originate in frontal cortex, rather than being calculated in the LC itself or elsewhere. Simultaneous recordings with high temporal resolution in-vivo will also help to delineate cause and effect in frontal cortex/LC interactions and will complement the accumulating data from human fMRI / pupillometry. Specific details of the above theory could then be tested; for example, in comparing the predictions for an LC driven purely by consideration of utility or estimation uncertainty, rather than by a state-prediction error as prescribed by Free Energy-based estimates. In-vivo recordings during the two tasks described here could also be examined for the characteristic patterns. For instance, in the pattern of LC firing predicted for the foraging task, the above modelling shows a sudden transition to a higher tonic level of activity during a change in the environmental statistics, and a much slower decay of activity occurring as rules stabilise. Triggering or blocking such patterns of firing during task performance would be particularly revealing regarding the proposed role of the LC.

Finally, we have not addressed the role of other neuromodulators that have related effects on behaviour. Whilst dopamine is explicitly included in Active Inference models as a precision parameter, other neuromodulators (e.g. serotonin) do not yet have a clear place within the model. Understanding the interplay between these systems will be crucial for placing LC activity in context – and will enable the explanatory power of Active Inference to be fully harnessed.

